# Maintenance of Bound or Independent Features in Visual Working Memory is Task-dependent

**DOI:** 10.1101/2021.04.12.439333

**Authors:** Ruoyi Cao, Yoni Pertzov, Zaifeng Gao, Mowei Shen, Leon Y. Deouell

## Abstract

Over the last decade, seemingly conflicting results were obtained regarding the question of whether features of an object are stored separately, or bound together, in visual working memory. Many of these studies are based on an implicit assumption about a default, or fixed, mode of working memory storage. In contrast, here we asked whether the anticipated memory probes used in a given experiment might determine the format in which information is maintained in working memory, consistent with its task-oriented function. In order to test this flexible maintenance hypothesis, we recorded EEG while subjects performed a delayed-match-to-sample task with different requirements and load. In Experiment 1, by contrasting conditions with and without the requirement of maintaining bound features, we found significant differences in EEG signals recorded in central-parietal channels while controlling for differences in visual stimulation. Such difference was confirmed by multivariate pattern analysis (MVPA). Moreover, behavior results, topographical distributions and cross-contrast MVPA decoding jointly suggested that this effect is not the same as task load effect. These results were confirmed in Experiment 2 with an independent group of subjects with the same paradigm, providing reliable evidence that the format of object maintenance in visual working memory could be flexibly shaped by the current task.

## Introduction

Our visual input is composed of a number of different visual features, including for example color, location, shape, orientation, and movement. In order to interact with the world efficiently, we need to not only correctly integrate different features into an object when viewing them, but also need to develop an internal representation of bound objects which remain accessible after the visual stimuli disappeared. How conjunctions of features are maintained has received a lot of attention in both the psychological and neuroscientific fields (Treisman & Gelade, 1980; Treisman, 1998;Schneegans & Bays, 2019a; Treccani, 2018). These studies largely focused on investigating on whether objects are stored in visual working memory (VWM) as bound objects or separated features.

Assessing the limits of VWM memory performance is a typical, if indirect, way to investigate this question. If features are separately stored in working memory, then performance should be limited by the number of features rather than the number of objects to be maintained. Otherwise, if bound objects rather than separate features are stored, then working memory performance should be limited by the number of objects regardless of the number of features. Following such reasoning, Luck and Vogel (1997) proposed a ‘strong object’ view since they observed that increasing numbers of features within objects did not impair performance in change-detection tasks. Moreover, invariance to the number of features was observed even when one object consisted of multiple values from the same feature dimension. Such a strong object hypothesis was supported by a study showing no difference in delay EEG activity between memorization of multiple and single color objects (Luria & Vogel, 2011).

However, other studies came to different conclusions. For example, Wheeler and Treisman (Wheeler & Treisman, 2002) observed that while features from different dimensions could be stored in parallel without a cost for working memory performance, adding features from the same feature dimension limited memory performance. They proposed the multiple-resources view, stating that VWM maintains features from different feature dimensions in parallel, while features from the same feature dimension compete for storage space. The multiple-resources view was later supported by other studies using either a similar (Delvenne & Bruyer, 2004) and different paradigms (Wang, 2017, Xu, 2002). For example, Wang et al (2017) found that for objects with two dimensions, the memory performance decreased as more feature values had to be remembered, but the other (fixed) dimension of that object was not affacted.

Yet another model suggests that VWM performance is limited by both the number of objects and the number of features. For example, Olson and Jiang (2002) found that when the number of objects was held constant, performance was better in the single-feature condition than in the multiple feature condition, suggesting that it was more difficult to store two features than one feature of an object. When the number of features was held constant, performance was better when features conjoined to form objects than when they were presented as isolated features. The importance of both the number of features and the number of items inspired the ‘weak-objects view’ of working memory limitation (Alvarez & Cavanagh, 2004; Hardman & Cowan, 2015).

Imaging studies have also supported both maintenance of separate features and of bound objects. For instance, both primate (Baizer, Ungerleider, & Desimone, 1991; Mishkin, Ungerleider, & Macko, 1983) and human studies (Courtney, Ungerleider, Keil, & Haxby, 1996; Smith et al., 1995) found that VWM for spatial location and item identities activated different regions of the brain. Other studies however found no evidence for distinct representations of spatial and non-spatial features (D’Esposito et al., 1998; Kravitz, Kriegeskorte, & Baker, 2010; Kravitz, Saleem, Baker, & Mishkin, 2011).

Debates about whether features are stored separately or bound are based on an implicit assumption that there is a default or fixed mode of VWM storage. However, such a universal mode cannot be taken for granted. Indeed, remarkable flexibility in prioritizing information in WM according to the task goal has been reported. For example, when non-human primates viewed the same to-be-remembered stimuli but were trained to expect different kinds of memory probes, delay activity in the prefrontal cortex showed different patterns (Rao, Rainer, & Miller, 1997). In an fMRI study in humans, participants were required to remember a face and a scene (Nobre, 2007). During the delay period, a cue was presented to inform which is to be tested later. Increased activity was observed in areas involved in face (fusiform gyrus) or scene (parahippocampal gyrus) processing, according to the cue. Indeed, VWM is not simply a passive representational state of visual input during a delay period, but is better conceived as a functional state bridging previous contexts and sensations to anticipated actions and outcomes (Myers, Stokes, & Nobre, 2017). Consequently, the anticipated memory probes (questions) used in a given experiment might actually determine the format in which objects will be maintained in VWM and the involved cognitive resources.

In order to test this flexible maintenance hypothesis, claiming that the brain stores separate features or bound objects according to the task goal, we recorded EEG while subjects performed a delayed-match-to-sample task with and without the requirement of maintaining binding between features. If there is a fixed mode for the system to store visual stimuli regardless of task requirement, we expected no systematic differences between these two conditions in the Event-Related Potential (ERP) nor any significant multivariate pattern separation (decoding) during the delay period, as the visual input to be maintained was identical. In contrast, if information maintained in VWM is adaptive to task goals, we expected to find such differences. In a control contrast, we tested whether a potential task-dependence effect is similar to the effect of memory load. Two experiments were conducted – the first one was exploratory, and the second confirmatory.

## Experiment 1

### Methods

#### Participants

Fifteen healthy volunteers from the Hebrew University of Jerusalem participated in the study. They were paid (40NIS/h, ∼$12) or given course credits for participation. All subjects had reportedly normal or corrected-to-normal sight and no psychiatric or neurological history. One subject did not finish the experiment and three were excluded from analysis due to noisy recordings. The remaining eleven subjects consisted of 6 males and 5 females (19–31 years old). The experiment was approved by the ethics committee of the Hebrew University of Jerusalem, and informed consents were obtained after the experimental procedures were explained to subjects.

#### Stimuli and apparatus

Subjects sat in a dimly lit room. The stimuli were presented with Psychotoolbox-3 (http://psychtoolbox.org/) implemented in Matlab 2018 on a ViewSonic G75f CRT (1024×768) monitor with a 100-Hz refresh rate. They appeared on a grey background at the center of the computer screen located 100 cm away from the subjects’ eyes.

Subjects performed a delayed match-to-sample test with 3 blocks of trials from 3 conditions (Figure 1): two-item-feature (F2), two-item-binding (B2), and four-item-binding (B4). In the B2 and F2 conditions the memory array consisted of 2 colored items randomly with identical irregular shape. Each item subtended a visual angle of 2.1°×2.1. The color of each item was randomly selected out of six highly distinguished colors, including red (RGB: 250,0, 0), green (0, 250, 0), yellow (250,250, 0), blue (0, 0, 250), violet(250,250,0) and white (0,0,0) (Figure 1), without repetition (i.e. each item had a unique color within a given array). The locations of the two selected items were randomly selected out of eight potential locations evenly distributed on an invisible circle with a diameter of 7.3°centered on the fixation cross. In the B4 condition, the memory array consisted of four items of different colors on four locations that were randomly selected from the six colors and eight potential locations respectively, without repetition.

**Figure 1.**
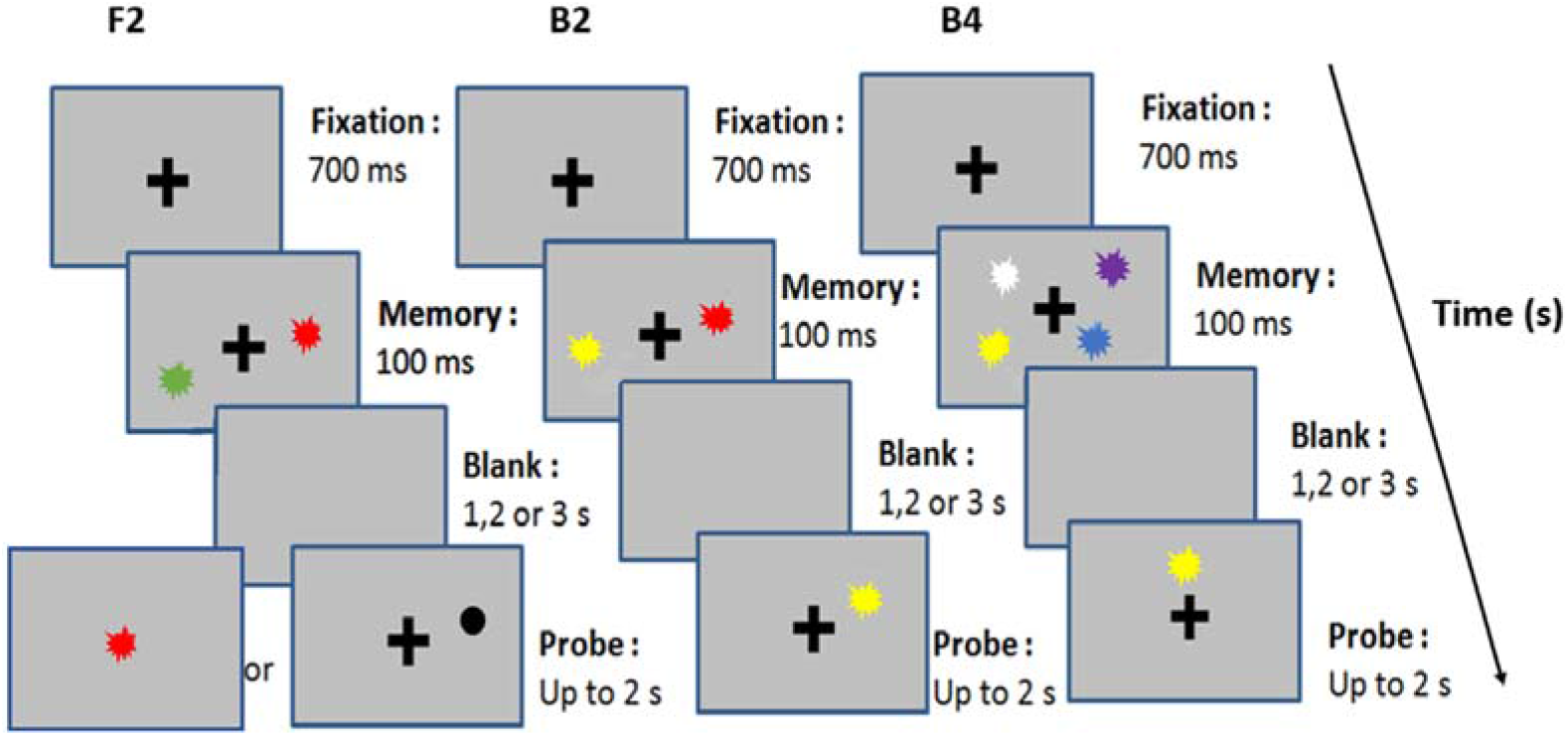
Illustration of an example trial in the three conditions: Left – Two-Item feature condition (F2) with Matched probe, middle – Two-Item binding condition (B2) with Mis-conjunction probe, right – Four-Item binding condition (B4) with New-feature probe.

Following a variable delay period of 1, 2 or 3 seconds, a single probe was shown on each trial. In the B2 and B4 condition, a probe could be one of three types: *Matched, New-Feature*, or *Mis-Conjunction*. A *Matched* probe had the same color and location as one of the items in the memory array. A *New-Feature* probe had either an old color but was located at a location not occupied in the memory array (“new”), or was at an location that was occupied (“old”), but had a color that was not used in the memory array. *Mis-conjunction probes* had an old color *and* an old location, both present in the memory array, but the conjunction between them was not used in the memory array. In the F2 condition, only Matched probes and New-Feature probes were used. A *Matched probe* in this condition was either presented at the center of the screen in an old color contained in the memory array, or a black item shown at an old location. Finally, a *New-feature probe* in this condition was a probe either presented at the center of the screen with a color that did not appear in the previous memory array or an item in black presented at a new location.

#### Experimental procedure

Each trial started with 700 msec a fixation cross presented at the center of the screen. Next, the memory array appeared for 100 msec. This memory array was followed by delay period of either 1,2 or 3 seconds blank screen, in equal proportions. Trials with the 3 delay durations were randomly mixed within a block. Following the delay period, a probe item was presented and subjects had to press a key to indicate if the probe item was an “old” or a “novel” item. The probe disappeared when the response was made (with a limit of 2 seconds), followed by the initial fixation of the next trial.

There were 3 conditions in the experiment, each consisting of 288 trials. In the B2 and B4 conditions, the delay period was followed by match probes in 144 trials, new feature probes in 48 trials, and *Mis-conjunction* probes in 96 trials. A probe was regarded as an old item only when the probe had both the color and the location of one of the items shown in the memory array. In the F2 condition, the delay period was followed by *match-feature* probes in 144 trials and the *new-feature* probes in the remaining 144 trials, and a probe was regarded old when either the color or the location of the probe had been shown in the memory array. Each condition began with 64 practice trials followed by 2 consecutive blocks of that condition, resulting in 6 blocks in total. Each block took about 15 mins, and subjects were instructed to take a break between blocks. The order of the three conditions was counterbalanced across participants.

As the memory arrays in B2 and F2 both included 2 items with 2 relevant features each, the contrast between B2 and F2 was intended to reveal the effect of retaining feature conjunctions in addition to individual feature values, keeping the visual stimulus (the memory array) identical. The contrast between B4 and B2 was conducted to reveal a load effect (maintaining 4 vs. 2 items).

All analyses and figures for the behavior results were made using JASP 0.8.5.1 (https://jasp-stats.org/). Response accuracy of 11 subjects was entered into a 3 (Delay duration: 1 vs 2 vs 3 sec delay) × 3 (Condition: F2 vs B2 vs B4) repeated-measures two-way ANOVA. Degrees of freedom were adjusted for violations of the assumption of sphericity with the Greenhouse–Geisser correction when necessary.

#### EEG recording and data analysis

EEG data were recorded using an Active 2 system (Biosemi, the Netherlands) from 64 active electrodes spread out across the scalp according to the extended 10–20 system with the addition of two mastoid electrodes and a nose electrode (https://www.biosemi.com/pics/cap_64_layout_medium.jpg). Horizontal electro-oculogram (EOG) was recorded from electrodes placed at the outer canthi of both eyes. Vertical EOG was recorded from electrodes placed above and below the left eye. The EEG was continuously sampled at 1024 Hz with an anti-aliasing low pass filter with a cutoff of 1/5 the sampling rate, and stored for off-line analysis. The data was referenced online to the Common Mode Sensor (CMS) which was placed in the space between POz, PO3, Pz, and P1.

Data preprocessing and analysis were done with the FieldTrip toolbox (version 20191213 http://www.fieldtriptoolbox.org/) implemented in Matlab 2018 (Mathworks, Natick, MA, USA). Preprocessing was applied to continuous data. During preprocessing, EEG and EOG signals were firstly filtered with Butterworth zero-phase (forward and reverse filter) bandpass filter of 0.1–180 Hz and then referenced to the nose channel. Extremely noisy or silent channels, which contributed more than 20% of all artifacts (Criteria: more than 100μV absolute difference between samples within segments of 100 ms; absolute amplitude > 100μV) were deleted. No more than 2 adjacent channels or 3 channels in a single subject were deleted, see details below. Next, data were re-referenced to an average of all remaining EEG electrodes. Ocular and muscular artifacts were removed from the EEG signal using the ICA method by manual selection of artifact components based on correlation with the EOG channels, power spectrum typical to muscle activity, and typical component scalp topographies. After ocular and muscle artifacts were removed, automatic artifact rejection was applied (http://www.fieldtriptoolbox.org/tutorial/automatic_artifact_rejection/). Time points larger than 12 standard deviations from the mean of the corresponding channel were marked, together with 200 msec before and after, so that the (subthreshold) beginning and end of an artifactual event will be likely accounted for. A visual inspection of the data followed in order to detect rare artifacts which were missed by the automatic procedure. Finally, previously deleted channels were recreated by mean interpolation of the neighboring electrodes (FC5, F6 were interpolated in subject 01, AFz in subjects 02, 04, and 06, AF8 and Iz in subjects 05, 07, and 09, and PO3 and CP1 in subject 08). Data were then down-sampled to 512 Hz, filtered with a Butterworth zero-phase lowpass filter at a cutoff at 20 Hz and parsed into 1800 msec segments starting 500 msec before the memory array onset, and averaged within each subject and condition. The average of 100 msec before the onset of stimuli over each trial was defined as baseline and subtracted from all data points of each segment.

There were two planned contrasts in our experiments: (a) the difference between F2 and B2, potentially revealing the effect of task on the ERPs. Since the two conditions differed only on whether the task was to retain features or bound objects given the same visual input, differences between these two conditions reflected effect of binding and was referred as “binding effect”. (b) the difference between B2 and B4, revealing the VWM “load effect” on the ERP, given the same task. To compare ERP amplitudes between conditions across all electrodes and time samples, cluster-based permutation tests (Maris & Oostenveld, 2007) were performed in Experiment 1, the exploratory phase. This approach allowed for a sensitive comparison between conditions at the level of spatiotemporal clusters without a predefined region of interest (ROI) and provided relevant correction for multiple comparisons with Monte Carlo based cluster-correction. The cluster-based permutation test included the following procedure: a) A paired sample t-test was applied to each time point and channel, resulting in one t-value for each data point. b) A threshold of p <.05 was applied to each time point on each channel, and a cluster was defined as the collection of above threshold data points, adjacent to each other either spatially or temporally. c) t-values were summed up within each cluster resulting in sum-t values. d) Repeating steps a-c for 1000 times on data while switching the condition labels within a randomly selected set of subjects for each iteration. Within each iteration, the largest positive and the smallest negative sum-t entered into the “positive” and the “negative” null distribution, respectively. e) The sum-t value of each cluster calculated in step c was compared with the two null distributions. Clusters with sum t-value larger than of 97.5% of the positive null distribution were defined as significant positive clusters, while clusters with the sum-t value smaller than 97.5% of the negative null distribution were defined significant negative clusters.

To examine whether the binding effect and the load effects had a shared topography, we used multivariate pattern analysis (MVPA). First, we tested for each subject whether we can predict the condition within each contrast (i.e. F2 vs B2 and B2 vs B4) based on the topographic patterns of brain activity at each time point. Next, we attempted to cross-decode, using the discriminating pattern found in one contrast to decode the conditions in the other contrasts. Finding significant cross-contrast decoding would suggest shared processes, possibly reflecting differences in task load in both contrasts.

MVPA was performed in MATLAB using the MPVA Light toolbox (https://github.com/treder/MVPA-Light). The classification was performed for each participant in a time-resolved manner. For each participant, trials in each condition were first randomly sub-sampled to the minimum number within all three conditions to attain a balanced set for training and testing. These data were then split into five equal parts. Then, a linear discriminant (LDA) classifier was trained for each time point from 200 msec before the stimuli onset to 1100 msec after the stimuli onset, using the voltage at 64 electrodes as features. For each fold, the classifier was trained on 80% of the data and tested on the remaining 20% of trials. To increase the signal-to-noise ratio, training and testing were done on averages over 4 trials, selected randomly with replacement within training and testing sets separately. The final results were averaged over the five folds. To attain decoding performance invariant to the decision boundary, Area Under Curve (AUC) was calculated for each subject. To reveal the generalization of these resulting classifiers over time, a temporal generalization matrix was computed for each subject, in which classifiers were trained on one-time point, and tested on all other time points (King & Dehaene, 2014). This provided detailed information about the sequence of processing stages engaged in the task. The above decoding procedures were applied on decoding two within contrast pairs (B2 vs B4 and F2 vs B2) to show the load effect and the binding effect respectively, and two across contrast pairs (Training on B2 vs B4, testing on F2 vs B2; Training on F2 vs B2, Testing on F2 vs B2) to show the independency (or dependency) of these two effects.

For each decoding analysis, a cluster-based permutation test was conducted to compare AUC with 0.5 with correction for multiple comparisons at the group level (Maris & Oostenveld, 2007). That is, the AUC of all subjects for each time point was first compared with 0.5 chance level with one-sample-test and then threshold at 0.01 significant level. Positive significant t-values of data points adjacent in time that survived the first level threshold defined clusters, and the t-values within each cluster was summed up to create t_sum_ values for each cluster (note that in the temporal generalization matrix, adjacent in time is defined as either adjacent in testing time or in training time). The t_sum_ values was compared with the distribution of the largest t_sum_ values in 1000 iterations generated by the same procedure, albeit with the chance level (AUC=0.5) and the true decoding performance switched in label for a random subsample of the subjects.

## Results

### Behavioral results

A 3 (Delay duration: 1 sec vs 2 sec vs 3 sec) × 3 (Condition: 2F vs 2B vs 4B) repeated-measures two-way ANOVA of response accuracy revealed a main effect of Condition, *F(2,20)* = 13.70, *p* < .001, *η2* =.58 (Figure 2). Follow-up pairwise comparison (with Bonferroni correction) showed a significantly lower response accuracy in condition B4 than both condition B2, mean difference *Mdiff* = - .095, *SE* = .03, *t* = -3.30, *p* =.011, *Cohen’d* =-.996, and condition F2, *Mdiff* = - .15, *SE* = .03, *t* = -5.18, *p* < .001, *Cohen’d* = -1.56, whereas no significant differences were found between B2 and F2 condition, *Mdiff* = - .054, *SE* = .03, *t* = - .56, *p* = .23. The main effect of Delay Duration was also significant, *F(2,20)* = 4.89, *p* = .002, *η2* =.328. Follow-up pairwise comparisons showed a marginally significant decrease of response accuracy from the 1 sec to the 3 sec delay, *Mdiff* = .03, *SE* = .01, *t* = 2.49, *p* = .065, *Cohen’d* = .75, and from the 2 sec to the 3 sec delay, *Mdiff* = - .026, *t* = 3.96, *SE* = .007, *p* = .006, *Cohen’d* = .75, but not from 1 to 2 sec delay, *Mdiff* = - .003, *SE* = .01, *t = -*.*26, p* = 1, reflecting some memory decrement following extended delays. The interaction between Delay Duration and the Condition was not significant, F*(4,40)* = .69, *p* = .60, *η2* =.07. These results indicated that the B4 condition, as expected, was more difficult than the other conditions, and that F2 and B2 conditions did not significantly differ in their overall exertion. Therefore, any results gained from comparing F2 and B2 conditions in EEG is less likely explained by differences in task difficulty.

**Figure 2:**
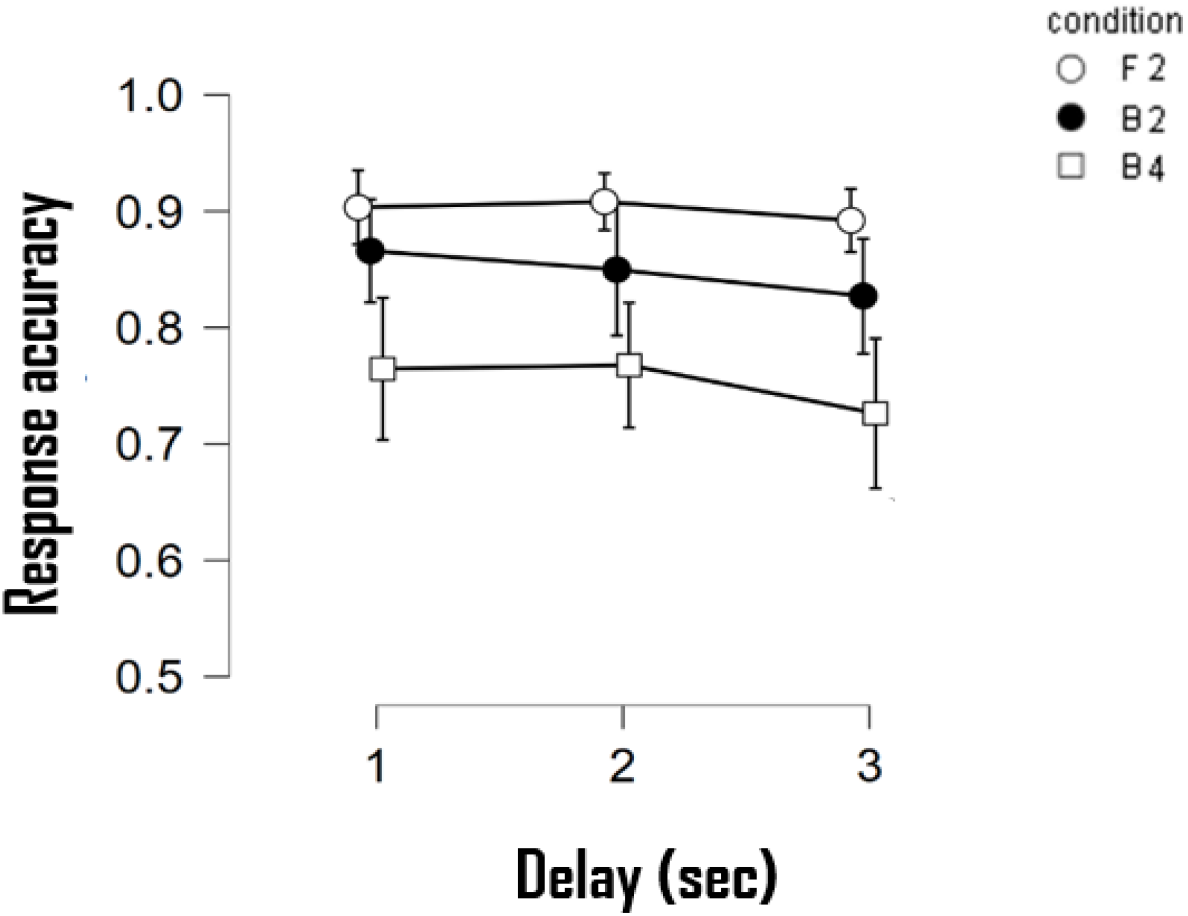
Percentage of correct responses on each condition following 1, 2, or 3sec delay between the memory and test. Error bar represents 95% confidence intervals.

### ERP results

#### Load effect: comparing B4 and B2 condition

We examined the effect of enhanced memory load by comparing the ERP amplitude in the B4 and B2 conditions up to the first second after the onset of the memory array (as this period was common to all 3 delays). Figure 3a shows that maintaining four-bound objects (B4 condition) evoked larger ERP amplitudes than maintaining two-bound objects (B2 condition) at right-frontal and middle-central electrodes. A spatiotemporal cluster-based permutation test showed that this significant cluster of difference (*p* < 0.05, cluster-corrected) extended from 394 msec to 847 msec after the onset of the memory array^1^. Figure 3b shows the grand average waveforms across subjects for 25 channels included in this significant positive cluster^2^.

**Figure 3 a:**
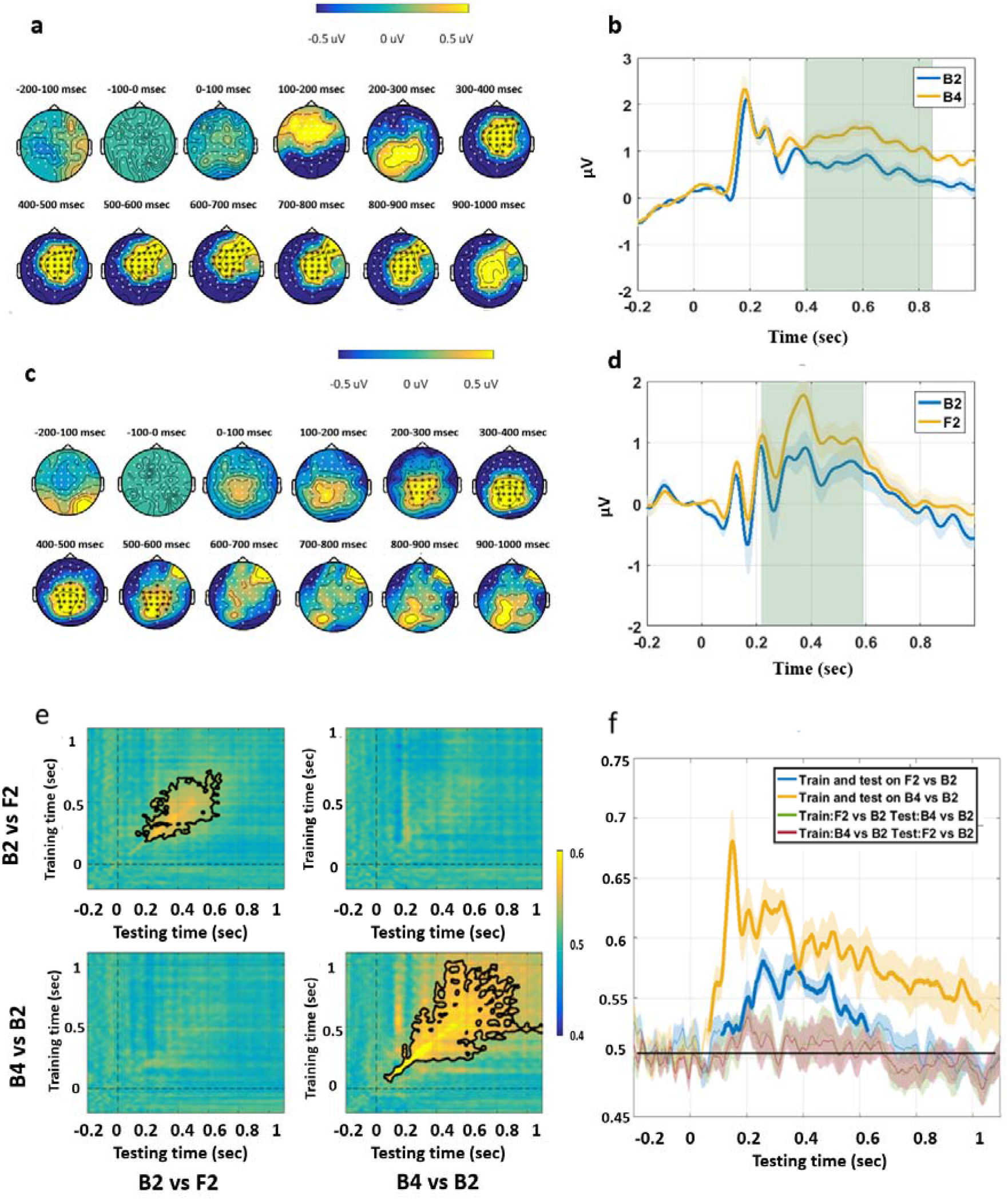
Difference in topographies of response amplitudes between B4 and B2 condition, in 100 msec bins, from 200 msec before until 1sec after the memory array onset. Yellow colors show higher positivity for B4 than B2. Black markers stand for channels included in significant clusters based on the cluster permutation procedure. **b:** ERPs elicited by the B4 and B2 conditions, averaged over subjects and channels included in the significant cluster. The light green rectangle highlights the temporal duration for the “Rectangle ROI” (see Figure 5a &Figure 5a). Shading around the waveforms represents standard error of the mean across subjects. 0 on the X-axis corresponds to the onset of memory array. **c**: Difference topographies between F2 and B2 conditions, in 100 ms bins from 200 ms before memory array onset. Yellow color indicates higher positivity in F2 conditions than in B2 condition while blue colors indicate the opposite. Black markers stand for channels included in significant clusters. **d:** As in **b**, ERPs elicited in the F2 and B2 conditions, averaged over subjects and channels in the significant cluster. **e:** Temporal generalization matrices. Training time is shown on the vertical axis and testing time on the horizontal axis. Significant clusters are indicated by black contours. **f:** Decoding performance (AUC) in Experiment 1, for classifiers trained and tested on the same time points (diagonals of the matrices in panel e). Thick lines indicate significant above chance decoding (cluster corrected, p<0.05). The shaded area represents the standard error across subjects.

#### Binding effect: comparing F2 and B2 condition

We compared ERP amplitudes when subjects were required to remember two items with (B2) and without (F2) the need to maintain the conjunction (binding) between color and location. As the memory array was identical in both conditions, a difference between the two conditions could not be due to differences in visual information. This contrast thus indicated whether the visual information had been maintained in a task-specific manner. Cluster-based permutation test showed that F2 elicited stronger positivity than B2 with a significant cluster in centro-parietal electrodes between 220 msec to 589 msec after the onset of the memory array (Figure 3c). Figure 3d shows the amplitude averaged across subjects and 20 channels included in this significant positive cluster.

### MPVA results

Multivariate analysis further revealed different responses between tasks. The temporal generalization matrices averaged over subjects are shown in Figure 3e. Cluster-based permutation test detected a significant cluster when training and testing on B4 vs B2 condition (Figure 3e bottom-right panel, *P* = 0.006, cluster-corrected). Figure 3f shows the diagonal of the temporal generalization matrix (Figure 3e), where testing and training were conducted on the same time points. Above chance decoding was found from 6 msec after stimulus onset until end of the maintenance period with the peak accuracy at 152 msec (*P* = 0.007 cluster corrected). Although the significant cluster started early after the stimuli onset, classifiers trained on early time points could not decode the later maintenance period above chance. Meanwhile, the later component starting around 200 msec after the stimuli onset generalized to the end of the maintenance period. This suggests that the significant contrast between B2 vs B4 found in the above ERP analysis reflected a VWM load effect, which is different from the early visual effect.

Training and testing on the same time point showed a significant decoding performance (AUC) between B2 and F2 from 111 msec to 630 msec with the peak accuracy at 259 msec (*P* < 0.001 cluster corrected Figure 3f), despite the identical visual presentation in these two conditions. This above chance decoding reflects a “binding effect”. The temporal generalization matrix indicated a relative stable temporal dynamic starting from around 200 msec after the stimuli onset and lasting until around 800 msec (Figure 3e Top-left panel, *P* = 0.068, cluster-corrected).

Next we asked whether we could decode B2 and F2 using a classifier trained on discriminating B2 and B4, and vice versa. Such cross-decoding would suggest shared spatial distributions between the “load effect” and the “binding effect”. However, despite the fact that we could decode B2 from F2 and B2 from B4 significantly above chance, we were not able to find a significant above chance cluster in the cross-decoding matrices, neither when training and testing at the same time points (diagonal) or across the generalization matrix. This was the case whether we trained on the B2 vs F2 contrast and tested on the B2 vs B4 contrast, or vice versa. This is consistent with the observation that the spatial distribution of the B2-B4 contrast was not identical to that of the B2-F2 contrast (compare Figures 3a and 3c).

Before discussing the results further, we note that the task-related differences were found through an exploratory method rather than a hypothesis-driven approach. Therefore, in experiment 2, we recruited a new group of subjects in an attempt to replicate this finding with the same paradigm.

## Experiment 2

### Method

We replicated Experiment 1 using the same experimental material and procedure.

#### Participants

Eighteen healthy volunteers from the Hebrew University of Jerusalem participated for either course credits or payment (40NIS/h). All subjects had normal or corrected-to-normal sight and reported no psychiatric or neurological history. Five subjects were excluded from analysis due to noisy recordings (subjects in which more than two neighboring channels were noisy were excluded). The final sample was composed of thirteen subjects (6 males, 7 females aged range 21-30) ^3^. Informed consents were obtained after the experimental procedures were explained to the subjects. The experiment was approved by the ethics committee of the Hebrew University of Jerusalem.

#### EEG recording and data analysis

Data preprocessing and analysis were the same as in Experiment 1. Briefly, preprocessing started with deleting extremely noisy channels. A Butterworth Bandpass filter of 0.1–180 Hz was applied. Ocular and muscular artifacts were extracted with ICA and removed. Then an automatic and manual artifact rejection was applied. Finally, previously deleted channels were recreated by mean interpolation of the neighboring electrodes (CP3 and P1 in subject 01; AF7, F7, P9 in subject 03; FC5 in subject 04; POz, Fp2 in subject 05; T7, TP8 and P9 in subject 06; F8 in subject 08, CP3, T7 in subject 09 ; T7, CP3 in subject 12) . After preprocessing, data were down-sampled to 512 Hz and were low-pass filtered with a Butterworth bandpass filter at a cutoff of 20 Hz, segmented, and corrected for the baseline.

In Experiment 2, the confirmatory phase, with a new group of subjects, we used pre-defined ROIs based on the significant spatio-temporal clusters found in Experiment 1. A “Rectangle ROI” for the contrast between two conditions was defined as all channels that were included in the significant cluster, spanning the interval from the earliest time to the latest time in that cluster across all channels. “Cloud ROI “included all channels within the significant clusters but including, for each channel, only the time points inside the significant cluster of that channel (Figure 5a &5d). One-tailed-paired-sample t-tests were then applied to compare amplitudes averaged over pre-defined ROIs (Rectangle ROI & Cloud ROI) between B2 and B4 conditions and F2 and B2 conditions respectively. The direction of the one-tailed test followed the expected effect as observed in Experiment 1.

## Results

### Behavior results

A 3 (Delay Duration: 1 sec vs 2 sec vs 3 sec) × 3 (Condition: F2 vs B2 vs B4) repeated-measures ANOVA revealed a main effect of Condition on response accuracy, *F*(2,24) = 27.38, *p* < .001, *η2 =* .*70*. Follow-up pairwise comparisons (all with Bonferroni corrections) showed a significantly lower response accuracy in condition B4 than both condition B2, *Mdiff* = - .15, *SE* = .023, *t* = -6.65, *p* < .001, *Cohen’d* = - 1.85, and condition F2, *Mdiff* = - .14, *SE* = .023, *t* = -6.13, *Cohen’d* = -1.70, *p* < .001, while no significant differences were found between B2 and F2, *Mdiff* = 001, *SE* = .023, *t* = 0.52, *Cohen’d* = 0.14, *p* = 1. These results are similar to the pattern of behavioral results in Experiment 1 and confirmed that B2 and F2 conditions were not significantly different in terms of task difficulty, while B4 condition was more tasking than the other two. As in Experiment 1, a main effect was found for Delay Duration, *F*(2,24) = 7.70, *p* = .003, *η2* = .39 (Figure 4). Follow-up pairwise comparison showed a significant decrease of response accuracy following 3 secs delay compared with 1 sec delay, *Mdiff* = - .032, *SE* = .01, *t* = -3.91, *p* = .009, *Cohen’d* = -1.09, while no significant difference in response accuracy was found between 1 sec delay and 2 sec delay, *Mdiff* = -.014, *SE* = .008, *t* = -1.74, *p* = .28, *Cohen’d* = -0.48, nor between 2 sec delay and 3 sec delay, *Mdiff* = - .02, *SE* = .008, *t* = -2.17, *p* = .12, *Cohen’d* = -.62. These results reflect forgetting of visual information over time. The interaction between Delay Duration and Condition was marginally significant, *F*(4,48) = 2.56, *p* = .05, *η2* =.18. Altogether, the behavior results confirmed the findings of Experiment 1.

**Figure 4:**
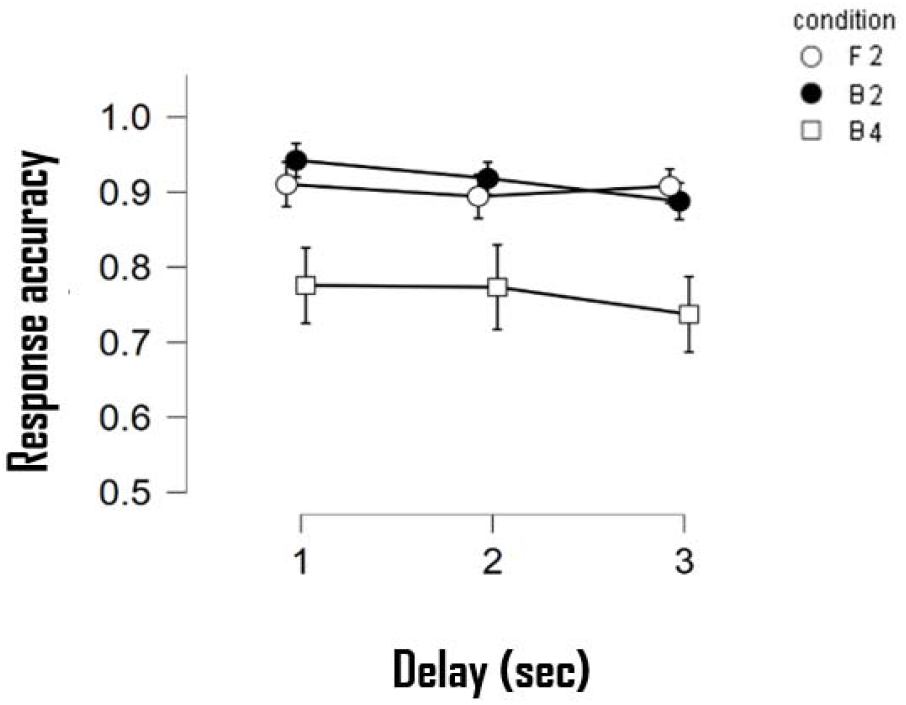
Percentage of correct responses on each condition following 1, 2, or 3sec delay between the memory and test in Experiment 2. Error bars represent 95% confidence interval across subjects.

### ERP results

#### Load effect: comparing B4 and B2 condition

Experiment 2 served as a confirmation stage following the exploratory stage of Experiment 1. Thus, the ROIs obtained from Experiment 1 were used to test for Condition Effect in Experiment 2, performed on a new group of subjects. Figure 5b shows the average ERP over subjects and channels included in the ROI. We averaged the amplitudes of the ERPs across the channels for the “Rectangle” ROI (Figure 5c dots in blue) including the channels consisting the significant cluster of Experiment 1, and all the time points within the cluster which were significant in any of the channels. The normality of the data was confirmed with Shapiro-Wilk test. A one-tailed paired sample t-test confirmed that, as in Experiment 1, the B4 condition *(M =* 1.13, *SE* = 0.24) evoked significantly higher positivity than the B2 condition *(M = 0*.*68, SE = 0*.*20), t(12) =* 3.14, *p* = .004, *Cohen’d =* 0.87. This direction of difference was found in eleven out of thirteen subjects (Figure 5c). A similar result (Figure 5c dots in red) was found using the “Cloud ROI”, which was the exact cluster found significant in Experiment 1, by channels and time-points, *t(12) = 3*.*06, p =*.*005, Cohen’s d = 0*.*85*. Thus, we successfully replicated the results showing that maintaining additional bound items in memory lead to larger positive response in frontal-right and middle-central channels within the time window ranging from ∼390 msec to ∼850 msec after the onset of the memory array.

**Figure 5:**
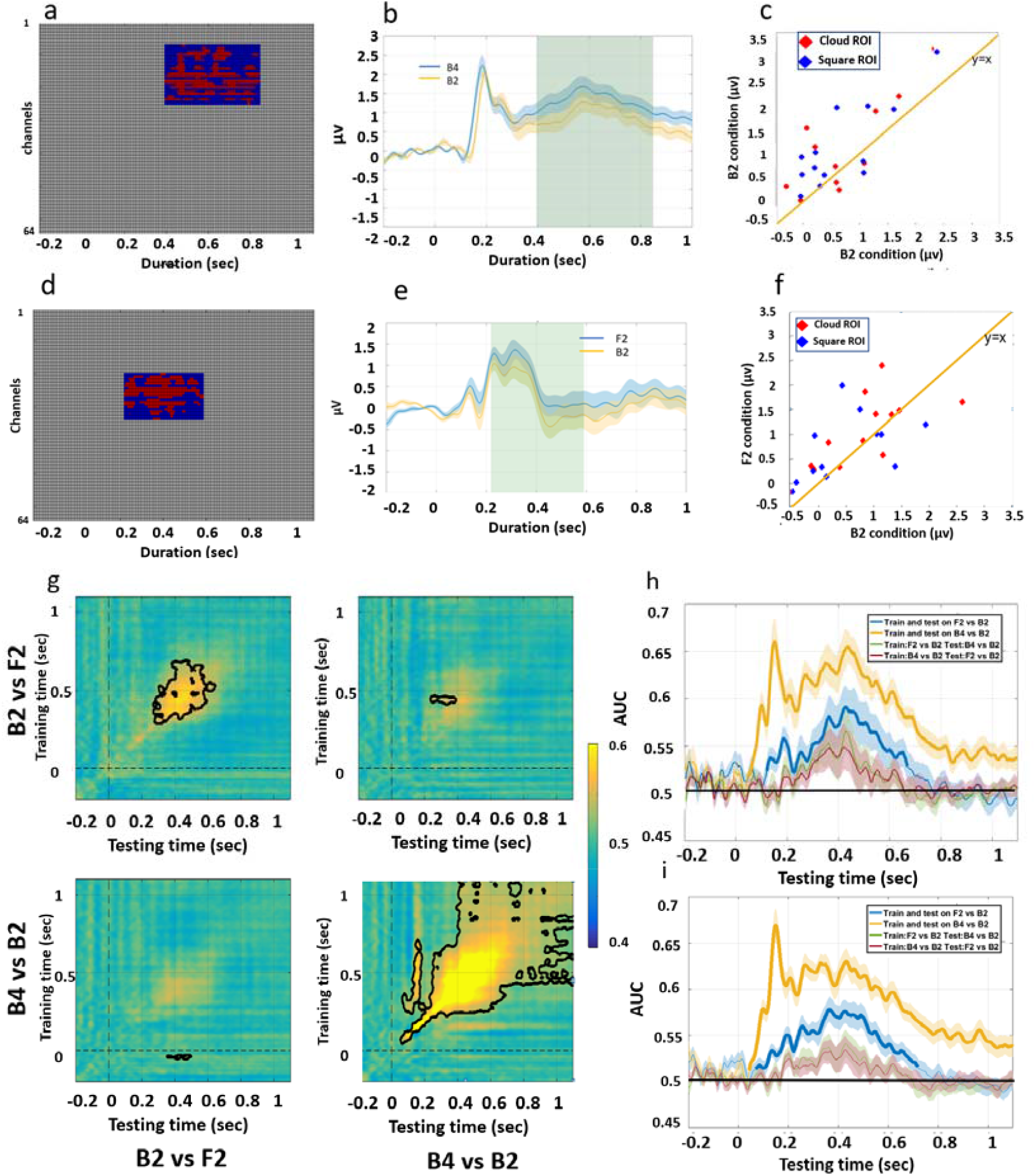
a:Matrix of channels x time, with red indicting time points and channels included in the significant cluster in the B2 vs B4 contrast in Experiment 1, representing the ‘cloud ROI’ for this contrast. The rectangle ROI (blue) applies the maximal temporal extent of the significant cluster to all the electrodes included in the cluster. From top to bottom, channels included in the significant clusters are AF4, AF8, F1, Fz, F2, F4, F6, F8, FC3, FC1, FCz, FC2, FC4, FC6, C3, C1, Cz, C2, C4, CP1, CPz, CP2, CP4, Pz, P2 b: ERPs elicited by the B4 and B2 conditions, averaged over subjects and channels included in the a priori cluster defined in Experiment 1. The light green rectangle highlights the temporal duration for the “Rectangle ROI”. Shading around the waveforms represents standard error of the mean across subjects. 0 on the X-axis corresponds to the onset of memory array. c: Blue dots and red dots represent the mean amplitudes elicited by B4 and B2 condition averaged over the ‘rectangle ROI’ and ‘cloud ROI’ for each subject respectively. d: As in a, for the F2 and B2 contrast. From top to bottom, channels included in the significant clusters are Cz, C3, C1, FCz, C2, C4, CP3, CP1, CPz, CP2, CP4, P3, P1, Pz, P2, P4, PO3, POz, PO4, Oz. e: As in b, ERPs elicited in the F2 and B2 conditions, averaged over subjects and channels in the ROI defined in Experiment 1. f: Blue dots and red dots represent the mean amplitudes elicited by B2 and F2 condition averaged over the ‘rectangle ROI’ and ‘cloud ROI’ for each subject respectively. g: Temporal generalization matrices from experiment 2. Training time is shown on the vertical axis and testing time on the horizontal axis. Significant clusters are indicated by black contours. h: Decoding performance (AUC) in Experiment 2 when classifiers were trained and tested on the same time points (diagonal of the generalization matrices in g). Thick lines indicate significantly above chance decoding (cluster corrected, p<0.05). The shaded area represents the standard error across subjects. AUC =0.5 is the chance level. i:The same as the h but with data in both experiments combined.

#### Binding effect: comparing F2 and B2 condition

The amplitude for F2 and B2 condition were averaged respectively within the spatio-temporal Rectangle ROI (Figure 5d). A One-tailed paired-sample t-test showed a strong trend for the F2 condition to evoke larger positivity (*M = 0*.*73, SE = 0*.*18)* than B2 condition (*M* = 0.40, *SE* = 0.22) within the Rectangle ROI. However this difference was not significant, *t(12) = 1*.*54, p =*.*08, Cohen’s d = 0*.*43*. Seven out of thirteen subjects exhibited larger averaged positivity in F2 than in B2 condition (Figure 5f, blue dots). A similar result was found when the Cloud ROI was applied (Figure 5f, red dots), *M = 1*.*056, SE = 0*.*691; t*(12) = 1.466, *p* =.086, *Cohen’s d = 0*.*41*.

### MPVA results

MPVA further revealed a “load effect” comparing the B2 and B4 contrast. Testing and training on the same time points achieved significant decoding accuracy from 48 msec after stimulus onset and lasted until the end of the maintenance period with the peak accuracy at 154 msec (*P* < 0.001 cluster corrected, Figure 3h). Cluster-based permutation test detected a significant cluster on the temporal generalization matrix (Figure 5g bottom-right panel, *P* = 0.005, cluster-corrected). As found in Experiment 1, a distinguishing pattern between B2 and B4 appeared early on, but only the later component starting around 200 msec generalized to the end of the maintenance period.

A “binding effect” was also found by MPVA. Training and testing on the same time point found a significant above-chance decoding performance (AUC) between B2 and F2 from 123 msec to 671 msec with the peak accuracy at 430 msec (*p* < 0.001 cluster-corrected). The temporal generalization matrix shows a stable significant cluster starting from around 200 msec after the stimuli onset and lasting until around 800 msec (Figure 5g Top-left panel, *p* = 0.069, cluster-corrected).

Next we asked whether “binding effect” and “load effect” are based on the same mechanism by attempting to discriminate B2 and F2 using a classifier trained on discriminating B2 and B4, and vice versa. Similar to the cross-contrast decoding result in Experiment 1, we were unable to decode across contrasts on either direction on Experiment 2, with the exception of a very short lived off-diagonal cluster shown in figure 5g (black contour in top right panel). No cross-contrast decoding was found when we pooled the subjects across experiments as well (Figure 5i).

To conclude, both the “load effect” and the “binding effect” was again showed by successful within-contrast decoding. Around chance cross-contrast decoding performances suggested that these two effects were distinct.

### Post-hoc analysis

#### ERP

The ERP over channels within the pre-defined ROI during the maintenance period reveals a trend for F2 condition to evoke larger positive ERP than the B2. A possible reason that the difference between F2 and B2 condition did not fully replicate is that the effect differs in the temporal domain across individuals. That is, a large difference between F2 and B2 conditions might appear soon after the offset of the memory array for some subjects, whereas for other subjects the difference between these two conditions takes longer time to develop. Such individual differences can be seen in Figure 6a, where the differences in amplitude between the F2 and B2 conditions for electrodes included in the pre-defined ROI are drawn for each subject, for every 200 msec bin from 200 msec to 1000 msec after memory array onset. It can be seen that the difference (F2-B2) was positive in most of the subjects and most of the time bins, although subjects varied as to the latency of the maximum difference. Therefore, we extended the temporal ROI into the whole maintenance period from 200 msec to 1000 msec after the array onset and segregated it into four 200 msec segments. We then averaged the amplitudes across the ROI channels for each segment. For each subject, we then selected the temporal bin that elicited the largest difference between F2 and B2, and compared the amplitudes of the F2 and B2 conditions across subjects with paired sample t-test. We compared the result to a distribution under the null hypothesis generated through a permutation procedure. In each iteration of this procedure we randomly permuted the labels of the B2 and F2 conditions and then applied the same procedure of segmentation and peak difference selection and compared the surrogate conditions with a paired sample t-test. Thus, each permutation contributed one value to the null distribution and this procedure was repeated 5000 times. The t-value of the empirical data (red vertical line in Figure 6b) was larger than 99.48 % of t-values in the permutated trials (Histogram in Figure 6b), which is unlikely to be obtained by chance. Thus, allowing temporal variance in the ROI of each individual leads to replication of the effect found in Experiment 1, showing that the F2 condition elicits larger positivity compared with B2 condition during the maintenance duration. Critically, we repeated the procedure, now looking for the most negative (F2<B2) difference between F2 and B2, but the result was far from significant in this case (Figure 6c).

**Figure 6:**
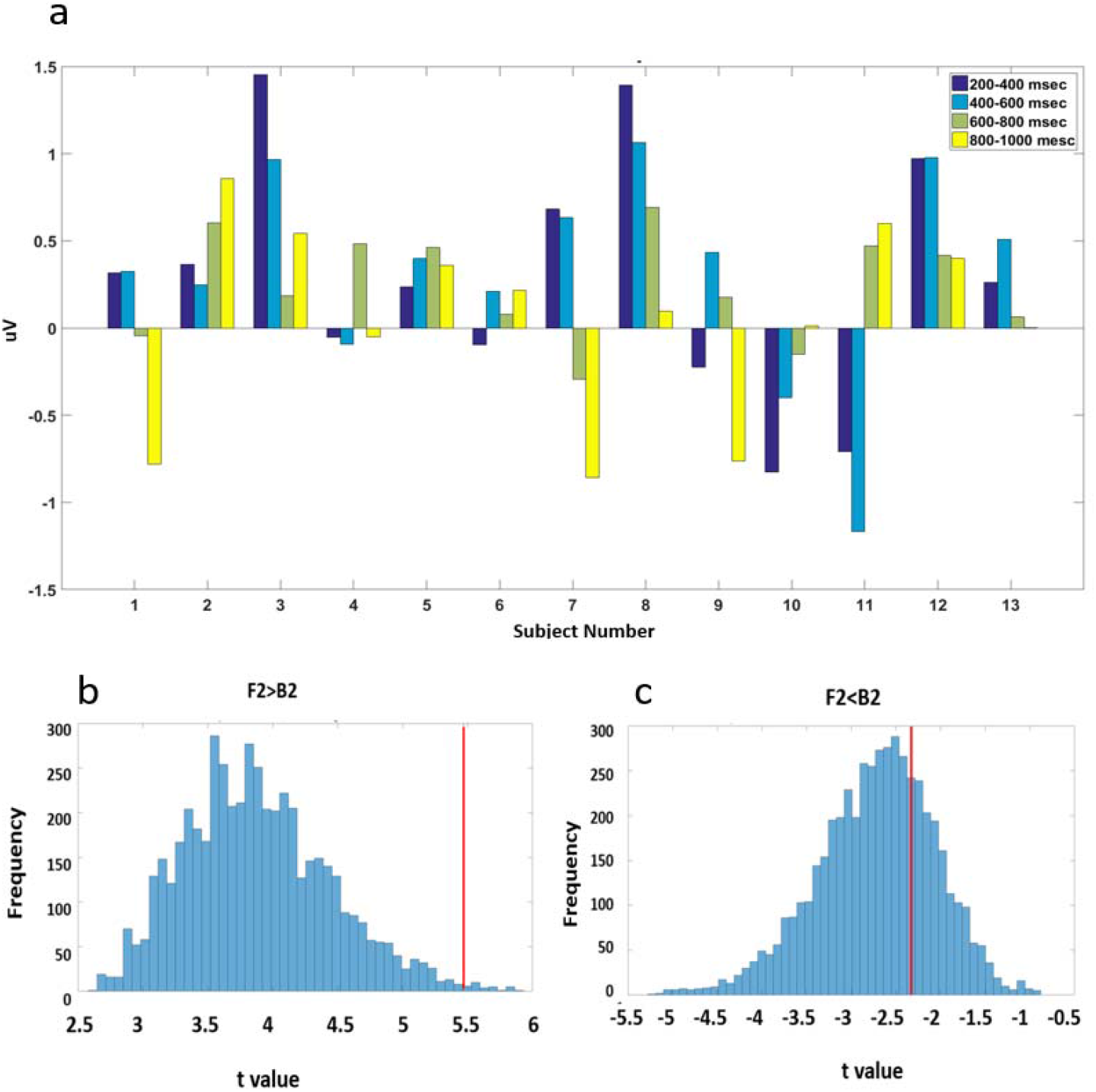
**a:** Amplitude differences between F2 condition and B2 condition (F2-B2) averaged across pre-defined spatial ROI for each subject within each 200 msec time window covering from 200 msec to 1000 msec after the memory array onset. **b**: The histograms, built by 5000 times randomly shuffling the label of B2 and F2 condition, represent the null distribution of t-values. The red lines represent the t-value of the empirical data. The t-value of maximum positive differences between F2 and B2 from 200 msec to1000 mesc after the onset of memory array is larger than 99.48 % results in null distribution. **c**: The t-value of maximum negative differences between F2 and B2 from 200 msec to1000 mesc after the onset of memory array is only smaller than 28.94% of the null distribution.

#### MVPA

If conditions in Experiment 2 could be successfully “decoded” from the classifier trained in Experiment 1, it would also support the notion that the same binding effect was found across the two experiments. Therefore, we trained group-based classifiers at each time point for Experiment 1 by pooling over all subjects and averaging each random 5 trials (sampled without replacement^4^) for each condition. The resulting classifiers were then applied to decode B2 from F2 condition for each subject in Experiment 2 at each time point. The testing set for each subject consisted of averages of 5 trials randomly selected with replacement. Classifiers trained in Experiment 1 were tested on every time point ranging from 200 msec before the stimuli onset until 1100 after the stimuli onset of each subjects in Experiment 2, resulting in a temporal generalization matrix of AUC for each subject. These results were then compared with chance level (*AUC=0*.*5*) at the group level by cluster based permutation test as used in the previous analysis. Classifiers trained during 310 msec to 538 msec after the memory array onset in Experiment 1 could decode successfully conditions in Experiment 2 from 314 msec to 851 msec (*p=0*.*049 clustered corrected, Figure 7*). This result is consistent with the observation that the “binding effect” found in the first group of subjects spreads into later times in the second group of subjects. Regardless of temporal window, this cross-experiment decoding analysis suggests that the topographical distribution characterizing the “binding effect” found in Experiment 1 was present also in Experiment 2.

**Figure 7:**
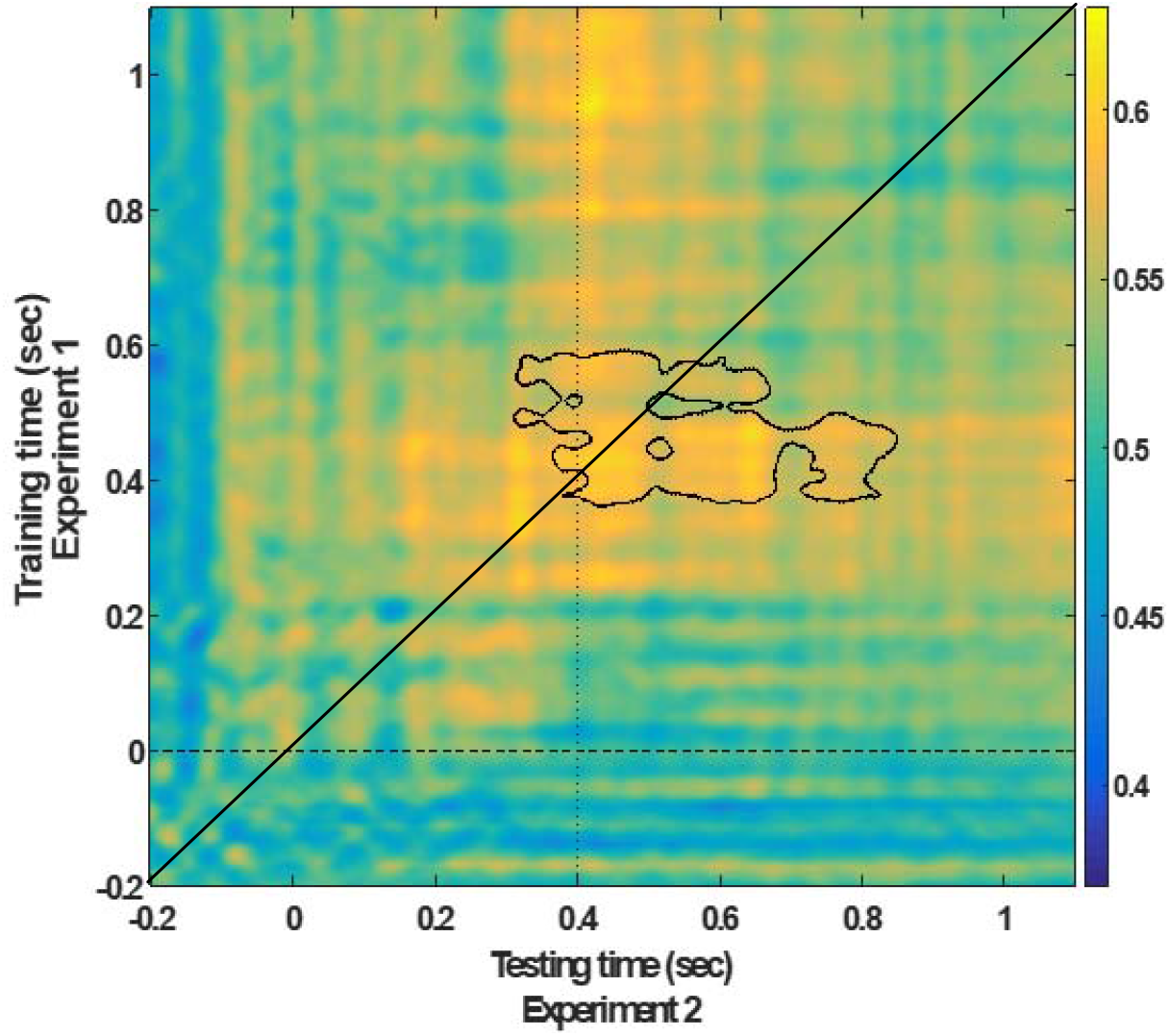
Temporal generalization matrix (average across subjects of Experiment 2) with training done on Experiment 1 (pooled across subjects) and testing on each subject of experiment 2. Training time is shown on the vertical axis and testing time on the horizontal axis. Significant clusters are indicated by black contours (cluster permutation).

## Discussion

In two experiments, we found significant differences in EEG signals between tasks requiring short term maintenance of visual features and maintenance of bound objects, even when the visual input was identical across conditions. This provides evidence of task-dependent maintenance processes and argues against the notion, implicitly embedded in previous literature, that a fixed default mode of item representation, either bound or not, is maintained in visual working memory. Furthermore, both topographic distribution and cross-contrast decoding results suggest that such differences found between tasks with and without binding requirements in VWM is not based on different levels of load involved in the task.

As expected, the ERP results reflected stronger central positivity in the more demanding B4 condition, compared to the B2 condition (Mecklinger & Pfeifer, 1996; Ruchkin & Johnson, 1990). Indeed, the behavioral results indicated that the B4 condition was more challenging than the other conditions. Considering the topographical distribution of this difference and the temporal dynamic, it is reasonable to take this increased positivity as a marker of memory load, but we cannot rule out a pure visual effect as the number of items on the display differed between B4 and the other conditions. Conversely, the B2 and F2 conditions consisted of identical arrays of two items, and lead to similar report accuracy. Still, a significant difference between the ERPs in these conditions was observed. Thus, the difference between the F2 and the B2 conditions was task-related rather than stimulus-related. The F2 condition elicited a higher central positivity and was more akin to the B4 (high load) condition than to the B2 condition in that respect. This pattern could reflect either (a) a larger number of units (i.e. 4 feature values vs. 2 bound objects) maintained during the delay period of F2 compared to B2 or (b) extra activity in the F2 condition when bound objects had to be decomposed into separate features to efficiently fulfill the task requirement. We note a more frontal topography for the B2 and B4 difference relative to the B2 and F2 difference, which might suggest different processes, favoring option (b) above. This is also confirmed by the cross-decoding results, suggesting that F2 vs. B2 effect is spatially different from the load effect. We also note that the MVPA analysis could discriminate F2 from B2 for a relatively limited duration of around 400 ms, which may fit more with a time-limited operation than with ongoing maintenance. However, these two explanations are not mutually exclusive and cannot be completely differentiated in the current study.

A previous visual search study (Berggren & Eimer, 2018) provided a piece of evidence that the brain represents information in a format compatible with the test demands. In this experiment, subjects answered whether the search display contained one of two target items held in working memory. When the search array included only one object with a target matching feature, targets and incorrect conjunction objects elicited identical N2pc component and sustained posterior contralateral negativity (SPCN). The N2pc is assumed to index a shift of spatial attention to the location of the potential target (Eimer, 1996; Kiss, Van, & Eimer, 2008), and the SPCN is assumed to index attentional activation of VWM representations of the potential target (Jolicoeur, Brisson, & Robitaille, 2008). These results suggest that in this condition only features were used as searching templates. In another condition, it was insufficient to detect a certain feature, as all objects had target-matching features, and both a target and an incorrect conjunction object could be present in the same display. In this case, the target evoked a larger N2pc than the incorrect conjunction objects, and only targets elicited SPCN components. Therefore, in this condition a bound object, rather than an isolated feature template, was used. Taken together, the results of this study are compatible with the possibility that VWM templates guiding attention in visual search were encoded or used flexibly, to effectively distinguish match and no-match probes, based on the task demand.

In our study, a color probe in the feature condition (F2) always appeared in the center (a location never used in the memory arrays) and a location probe was always colored black (a color never used in the memory arrays). Thus, the matched-probes in F2 (feature) condition matched only one feature dimension of the stimuli in the memory array, whereas the other feature of the probe was fixed across trials, with a value that never appeared in the memory array. In this condition, it is more efficient to maintain feature values which are not yoked to other feature dimensions. Since the memory array contained two colors and two locations, 4 different feature values had to be maintained. The matched-probes in the B2 (binding) condition, on the other hand, matched an item from the memory array in both feature dimensions. In this case, retaining bound objects allows a direct comparison between the representation in the VWM and the upcoming probe, and may lead to more efficient performance.

Task-specific encoding and retention of visual stimuli have been observed by other studies addressing VWM. Regarding the integration of object parts, a recent study (McCants, Katus, & Eimer, 2020) using CDA showed that objects were represented separately by their parts when parts appeared as probes, whereas a single compound object was maintained when objects were used as probes. Addressing the question of feature binding, one study found that memory for different features can be differentially affected by retro-cues indicating which feature dimension would be tested (Park, Sy, Hong, & Tong, 2017). Woodman and Vogel (2008) suggested individuals can control which features of an object are selectively stored in working memory. In this study, a lower contralateral delay activity (CDA), a marker of memory load, was observed when only color was task-relevant compared to the case both color and orientation were relevant. Notably, in the feature condition in those paradigms only one feature had to be maintained, allowing the subjects to ignore the other feature altogether, whereas in the current experiments, both features had to be maintained in both conditions, albeit in different formats. Our findings suggest therefore that given the same low level features to be represented in VWM, the task goal determines whether they are kept bound as items or separately as features.

The finding that objects are retained either as separate features or as bound items based on task requirements may explain some of the seemingly conflicting results in the literature, although not all of them. This is based on the idea that although two tasks are similar in their explicit instructions, subtle differences in the relationship between the memory array and the probe might affect the way that the memory array is maintained in memory. For example, Luck and Vogel (1997; extended in Vogel, Woodman, & Luck, 2001) observed that the ability to detect a change in a stimulus array of a given set size is the same regardless of the number of features. In contrast, Wheeler and Treisman (2002) found memory capacity is determined merely by the number of total features when two features from the dimension were combined into one object. In Luck and Vogel (1997), each item presented in the memory array contained features randomly selected from all possible feature values with replacement (e.g., more than one item could be red) and the non-match probe could include an erroneous conjunction of features that appeared in the memory array. In contrast, in the study by Wheeler and Treisman (2002, Experiments 1 and 2), items on the memory array were generated by selecting from possible feature values without replacement (e.g. only one item could be red). Moreover, to generate a non-matched probe, the probe included a feature that had not been used by any item in the memory array. This design, where no replacement was allowed when assigning feature value to a single memory array, can be also found on other studies claiming that increased features to be remembered for each object impaired change detection (Oberauer & Eichenberger, 2013). Therefore, in the study by Luck and Vogel (1997), bound objects had to be maintained to detect a non-match probe, while in studies showing contradictory observations, remembering a list of all features in the memory array was sufficient to distinguish between matched and non-match probe. The different results regarding representation format might therefore reflect the fact that such format is flexibly shaped by task goal. The current findings, therefore, draw attention to subtle details between experimental designs which could have an impact on the pattern of information that is maintained in memory.

Whereas the electrophysiological difference between the B2 and B4 condition was accompanied by a difference in task accuracy, the same difference between B2 and F2 conditions did not result in an observable change in accuracy. This highlights a dissociation between the activity measured continuously during the maintenance period, and the decision recorded after the presentation of the probe. Taking the results together, one can speculate that more units held in VWM are reflected by enhanced positivity (and possibly an enhancement of the CDA when stimuli are lateralized), whereas diminished accuracy reflects increase in difficulty of comparing the probe with what was retained in working memory. For example, Alvarez and Cavanagh (2007) showed that VWM capacity is inversely correlated with item complexity. However, Awh et al (2007) showed that the sample-test similarity is highly correlated with item complexity, and that the item complexity effect on memory performance was not observed when the sample-test similarity was low. This suggests that the retrieval process, rather than the maintenance stage, limits memory performance. Further studies could test this hypothesis more directly.

It is worth noting that we have used location-to-color binding in our study, and this kind of binding has some unique aspects that might not generalize to other types of binding. There is some evidence suggesting that objects are automatically encoded together with location. For example, some studies showed the task-irrelevant location of a stimulus can also be directly decoded from EEG data during the delay period of a VWM task (Elsley & Parmentier, 2015; Foster, Bsales, Jaffe, & Awh, 2017; Olson & Marshuetz, 2005). EEG studies show that spatial attention can be drawn to items in VWM even when the cue is non-spatial and the location is entirely irrelevant for the task (Eimer & Kiss, 2010; Kuo, Rao, Lepsien, & Nobre, 2009). There are studies supporting an even stronger claim that non-spatial features binding is mediated by binding to locations (Schneegans & Bays, 2017; Treisman & Gelade, 1980; Pertzov & Husain, 2014), perhaps due to the fact that most neurons selective to visual features also exhibit some spatial selectivity (Schneegans and Bays, 2019). Further studies are needed to test whether the results in the current test generalize to cases in which objects are defined by two non-spatial features.

To conclude, our study supports the view that VWM should not be construed as a passive storage of environmental input, but a dynamic system actively engaging in the task and preparing to carry out an effective response. Specifically, it provides a novel piece of evidence that the format of objects represented in the working memory, either as bound objects that integrate different feature dimensions or as separated features, is task-dependent. The finding of flexible maintenance of features and objects calls for a reconsideration on how to conceptualize and approach the binding problem in VWM.

## Acknowledgements

The study was supported by a China-Israel cooperative *scientific* research provided by the Israel ministry of Science, Technology and Space to Leon Y. Deouell and Yoni Pertzov, and Ministry of Science and Technology of the People’s Republic of China (2016YFE0130400) to Mowei Shen. We thank Yongdi Zhou for helping with design and discussion of the present study. We thank Naomi Revel and Geffen Markusfeld for helping with running experiments.

## Footnotes

1 Note that the cluster analysis does not ensure that the effect in any single point in the cluster is independently significant (Sassenhagen & Draschkow, 2019)

2 Note that a negative cluster on the left-temporal region is occasionally significant (P<.05), but it was not stable across multiple run in permutation.

3 Given the effect size of the B2 and F2 contrast in experiment 1 using the “cloud” ROI (the smallest effect size in all possible contrasts), this number of subjects gave power of 96% with 0.05 alpha level in Experiment 2.

4 Unlike the single subject decoding, due to the large number of trials, averaged data was made without replacement to reduce computational load.

